# *De novo* cancer mutations frequently associate with recurrent chromosomal abnormalities during long-term human pluripotent stem cell culture

**DOI:** 10.1101/2024.07.01.601640

**Authors:** Diana Al Delbany, Manjusha S Ghosh, Nusa Krivec, Anfien Huygebaert, Marius Regin, Chi Mai Duong, Yingnan Lei, Karen Sermon, Catharina Olsen, Claudia Spits

**Affiliations:** Research Group Genetics, Reproduction and Development, Vrije Universiteit Brussel, Laarbeeklaan 103, 1090, Jette, Belgium; Vrije Universiteit Brussel (VUB), Universitair Ziekenhuis Brussel (UZ Brussel), Clinical Sciences, research group Genetics, Reproduction and Development, Centre for Medical Genetics, Laarbeeklaan 101, 1090 Jette, Belgium; Brussels Interuniversity Genomics High Throughput Core (BRIGHTcore), Vrije Universiteit Brussel (VUB), Université Libre de Bruxelles (ULB), Laarbeeklaan 101, 1090 Brussels, Belgium; Interuniversity Institute of Bioinformatics in Brussels, Université Libre de Bruxelles (ULB) - Vrije Universiteit Brussel (VUB), Brussels, Belgium

**Keywords:** human pluripotent stem cells, genetic instability, copy number variation, single nucleotide variants, cancer-related genes

## Abstract

Human pluripotent stem cells (hPSC) are pivotal in regenerative medicine, yet their *in vitro* expansion often leads to genetic abnormalities, raising concerns about their safety in clinical applications. This study analyzed ten human embryonic stem cell lines across multiple passages to elucidate the dynamics of chromosomal abnormalities and single nucleotide variants (SNVs) in 380 cancer-related genes. Prolonged *in vitro* culture resulted in 80% of the lines acquiring gains of chromosomes 20q or 1q, both known for conferring *in vitro* growth advantage. 70% of lines also acquired other copy number variants (CNVs) outside the recurrent set. Additionally, we detected 122 SNVs in 88 genes, with all lines acquiring at least one *de novo* SNV during culture. Our findings show higher loads of both CNVs and SNVs at later passages which are due to the cumulative acquisition of mutations over a longer time in culture and not to an increased rate of mutagenesis over time. Importantly, we observed that SNVs and rare CNVs follow the acquisition of chromosomal gains in 1q and 20q, while most of the low-passage and genetically balanced samples were devoid cancer-associated mutations. This suggests that the recurrent chromosomal abnormalities are the potential drivers for the acquisition of other mutations.

## INTRODUCTION

hPSCs are an invaluable resource for regenerative medicine, with over 100 clinical trials currently ongoing (Kobold et al., 2023). At low passages, most hPSC lines maintain a normal diploid karyotype. However, during *in vitro* expansion, hPSCs frequently acquire genetic aberrations, commonly involving full or segmental gains of chromosomes 1, 12, 17, and 20 (Amps et al., 2011; Baker et al., 2016). These genetic aberrations are reminiscent of those seen in cancers, raising concerns about the potential for oncogenic transformation in transplanted hPSC-derived cell products (Andrews et al., 2022). Four studies have addressed this concern by investigating the incidence of point mutations in cancer-related genes (Avior et al., 2021; Lezmi et al., 2024; Merkle et al., 2017, 2022), and confirmed that hPSCs carry potentially deleterious variants in these genes, with *TP53* being the most commonly mutated. While these studies provided valuable insights into the mutational landscape of hPSCs, they could not reliably determine the timing of the appearance of these variants or their association with other mutational events such as the acquisition of CNVs. In this work, we close this gap in knowledge by studying multiple passages of ten hESC lines using simultaneous targeted gene sequencing with a panel of 380 cancer-associated genes and CNV analysis via shallow whole-genome sequencing to investigate the timing and association of these genetic variants. This approach provides insight into the temporal evolution of genetic changes within each hESC line.

## RESULTS

Figure 1A shows an overview of all the gains and losses identified, Figure 2 shows the genetic variants identified per cell line and passage, and a complete list of CNV breakpoints is provided in Supplementary Table 1. Of the 10 cell lines, only VUBe005 showed a balanced genetic content at all tested passages (P39, P75 and P89). One line (VUBe026) already carried CNVs at the earliest passage tested (P11), and the other 9 lines acquired different chromosomal abnormalities during extended culture. Overall, gains were more common than losses (39 gains vs. 8 losses). We found that 80% of our lines (8/10 lines, 13/33 samples) acquired a gain of chromosome 20q and 60% (6/10 lines, 8/33 samples) a gain of 1q, both well-known highly recurrent chromosomal abnormalities that confer a growth advantage to hPSCs *in vitro* (Avery et al., 2013; Krivec et al., 2023; Nguyen et al., 2014; Stavish et al., 2023). All gains on 20q included the driver gene *BCL2L1*, had a common proximal breakpoint and varying distal breakpoints, with sizes ranging 1.075 Mb to 46.925 Mb. The proximal breakpoints for the gains of 1q were specific to each line and were telomeric, their sizes ranging 0.725 Mb to 103.7 Mb and spanning the driver gene *MDM4*. These findings fully align with previous reports on these recurrent abnormalities (Amps et al., 2011; Halliwell et al., 2021; Merkle et al., 2022). Two lines also carried losses of 18q, a recurrent but less common chromosomal abnormality in hPSCs (Amps et al., 2011; Baker et al., 2016; Spits et al., 2008; Stavish et al., 2023). Further, we found an array of other abnormalities, including duplications of 1p13.2, 1q21.3, 3q26.22q27.3, 7p22.3pter, 9p24.3p13.3, 15q26.1q26.2, Xp11.3p11.23 and losses of 2q37.1, 6p21.33, 16p12.2p12.1 and 20p13, none of them being typically observed aneuploidies in hPSCs. These genetic changes do not appear to contain any genes with functions that would make them obvious driver genes for an *in vitro* selective advantage (listed in supplementary table 1). In all but one case, they appeared together with the recurrent genetic changes, suggesting that they may be passenger events. The analysis of datapoints of all lines combined shows that the later the sampling, the higher the load of acquired CNVs (p<0.001, Poisson Loglinear Regression, Figure 1B). This association is not retained when considering the absolute passage number (p=0.07, Figure 1C), but if the two latest passages are removed as outliers (passages 285 and 353), the association becomes statistically significant (p=0.032). Conversely, later passages did not have a higher risk of acquiring a *de novo* CNV as such (p=0.102, Binary Logistic Regression, Data not shown). This suggests that the higher loads of CNVs seen at later passages are due to the cumulative acquisition of mutations over a longer time in culture and not to an increased rate of mutagenesis over time.

**Figure 1.**
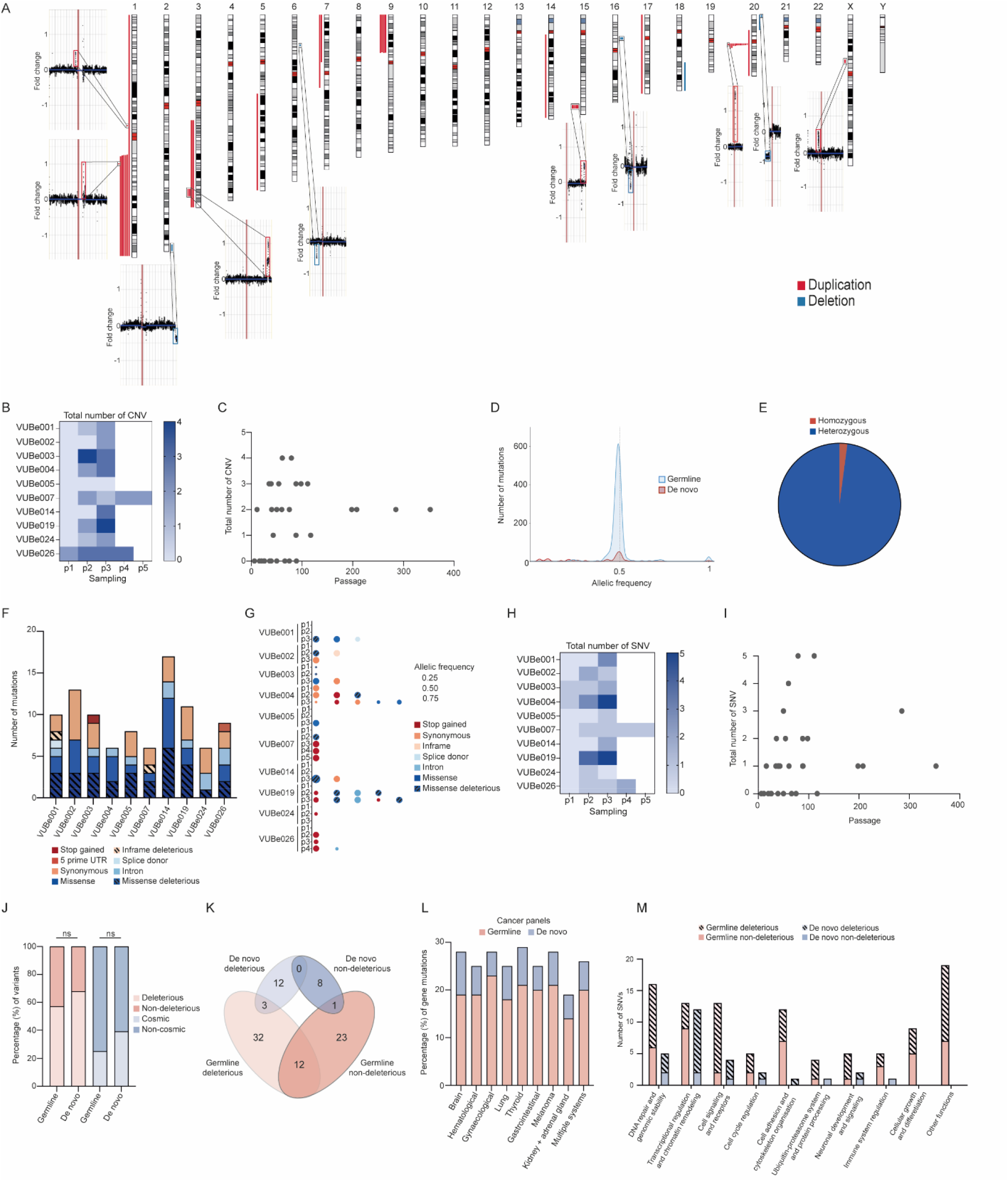
CNVs and point mutations identified in hESC samples from 10 different lines at multiple passage. **A**. Ideogram showing the location of all gains (red) and losses (blue) found in the 10 hESC lines, as well as the shallow sequencing results of samples in which we found unique CNVs, such as gains on chromosomes 1, 3, 7, 9, 15, X, and losses on chromosomes 2, 6, 16 and 20 that are not recurrent in hPSCs. **B**. Heatmap of the total CNV across different passages for each cell line. Each column (P1, P2, P3, P4, P5) represents the sequential sampling points (first, second, third, fourth, and fifth passages tested, respectively). **C**. Total number of CNV vs culture period *in vitro* across all the hESCs lines. **D**. Histogram representing the distribution of allelic frequencies of all mutations grouped by mutation origin (germline, and *de novo*). **E**. Distribution of detected germline mutations by zygosity. The pie chart illustrates the proportion of homozygous (red) vs heterozygous (blue) mutations identified in the samples. **F**. The bars represent the distribution of mutation categories for germline SNPs and the number of events detected in each hESC lines. **G**. Overview of *de novo* SNPs, the type of mutations detected in each passage of each hESC line, and the read frequency of each variant. Large bubble: high allelic frequency; average bubble: intermediate allelic frequency; small bubble: low allelic frequency; no bubble, no variant. **H**. The heatmap shows the incidence and the number of de novo mutations found in hESC lines depending on their culture period *in vitro* (Poisson regression **P=0.003). **I**. Total number of SNV vs culture period *in vitro* across all the hESCs lines. **J**. Proportion of deleterious and non-deleterious variants (%) found among both germline and *de novo variants* and whether they see are reported in the COSMIC database. **K**. Diagram showing the number of genes with germline (pink) or *de novo* (blue) mutations, and their pathogenic effect. **L**. The distribution of gene mutations (%) across different cancer panels, categorized into germline (pink) and *de novo* mutations (blue). The y-axis represents the percentage of gene mutations, while the x-axis lists the various cancer types. **M**. The bar graph illustrates the number of SNVs across various functional categories, differentiating between germline and *de novo* mutations, as well as their deleterious and non-deleterious effects.

**Figure 2.**
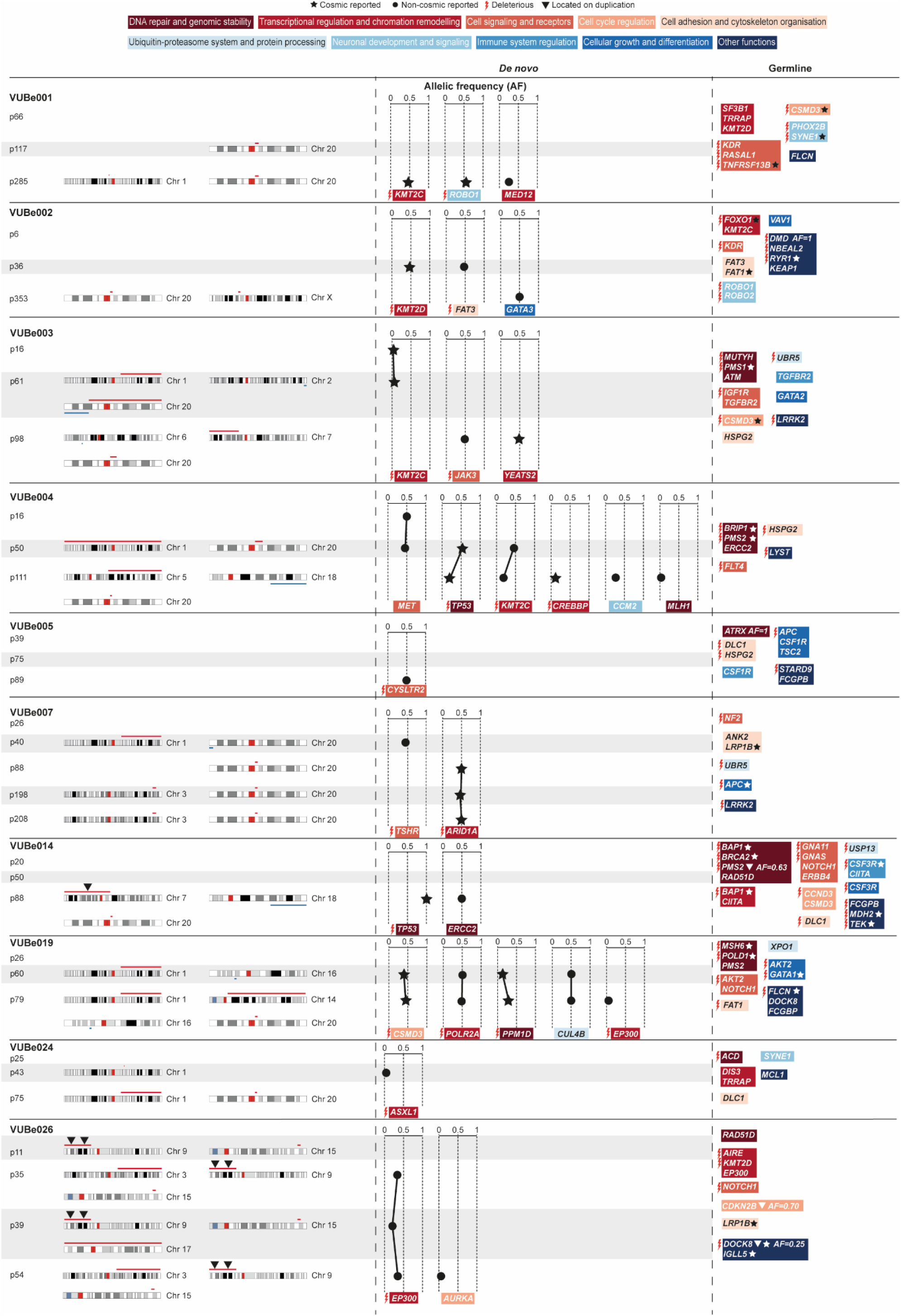
Distribution of chromosomal abnormalities and point mutations in cancer related genes, with a focus on their functional impact in hESC lines. Chromosomal abnormalities acquired at a specifi passage are shown for each hESC line, with their locations (left section). Point mutations found are shown in the middle (*de novo*) and right (germline) sections of the figure. Each mutation is annotated with the affected gene, the nature of the mutation, and its functional category, as shown by the function colour key at the top of the figure. The allelic frequency (AF, scale 0 to 1) of *de novo* variations is indicated for each gene. Mutations reported in COSMIC database are marked with (⍰), mutations not reported in the database are marked with (•). Mutations that are expected to be harmful (deleterious) are denoted ( ). Mutations found in duplicated regions of the genome are marked with (⍰).

With regards to the results of the cancer-associated gene sequencing, we identified a total of 122 single nucleotide variants (SNVs) in 88 genes across the different passages of the 10 lines (listed in supplementary table 2 and overview in Figure 2). While all 122 SNVs were different, 25 of the 88 genes were found to carry two or more SNVs. Since we sequenced samples of multiple passages of the same line, we could determine which of the SNVs were there from the onset of the establishment of the cell line (N=96, assumed to be germline SNVs) and which appeared *de novo* during cell culture (N=28). The allelic frequency of 95.8% of the detected germline SNVs (92 of 96) was around 0.5, as expected for heterozygous alleles, two of the germline SNVs were homozygous, and three SNVs had allelic frequencies of 0.25, 0.7 and 0.63 respectively (Figure 1D). *De novo* SNVs had a peak in allelic frequency at around 0.5, but also frequently appeared with lower allelic frequencies, and once as homozygous (Figure 1E). All our hESC lines carried germline SNVs, ranging from 6 to 17 SNVs per line, with a similar distribution in type of functional impact across lines (Figure 1F). From the 96 germline SNVs, 52 were missense SNVs, 29 of which predicted to be deleterious mutations by SIFT prediction tool (https://sift.bii.a-star.edu.sg/). We also found 28 synonymous mutations, 1 stop-gain mutation, 5 SNVs in a splice site, 1 in the 5’ untranslated region, 2 in-frame deletions and 7 mutations in introns.

All hESC lines acquired at least one *de novo* SNV, with a maximum of 5 SNVs in the later passages (Figure 1G and Figure 2). Only two out of the ten lines showed *de novo* SNVs at the earliest passage tested (VUBe003 and VUBe004). Overall, the SNVs were predominantly found in samples that had been extensively passaged (mid and late passages), indicating that hESCs are more likely to acquire *de novo* variants during prolonged *in vitro* culture (p=0.002, Poisson Loglinear regression Figure 1H). The absolute passage number did not significantly associate to the number of *de novo* SNVs, even after removal of the outlier passage numbers (p=0.101, Poisson LogLinear Regression, Figure 1I). Similarly, the rate of *de novo* mutagenesis does not increase with time in culture (p=0.771 for passage number, p=0.152 for sample rank, Binary Logistic Regression, Data not shown), suggesting that the SNV mutation rate stays constant with time in culture. Overall, of the 40 *de novo* SNVs, 16 were missense mutations, of which 13 predicted to be deleterious mutations, 7 synonymous changes, 12 stop-gain mutations, 1 in splice site region variant, 1 in-frame deletions and 3 SNVs in introns. None of the *de novo* SNVs we found progressively increased in allelic frequency across the passages we tested. While some SNVs appeared at low frequency in the earlier passages, their frequency over extended passaging did not change, suggesting that the cell fraction carrying the variant did not increase. Other variants decreased in frequency, or they disappeared entirely in subsequent passages of the same line (Figure 2).

We categorized the mutations based on their potential deleterious effect, including stop gains and missense variants, and assessed their clinical relevance by checking if they had been previously reported in the Catalog of Somatic Mutations in Cancer (COSMIC: https://cancer.sanger.ac.uk/cosmic), irrespective of their tier. Remarkably, most of the identified germline and *de novo* variants were potentially deleterious, with no statistically significant differences between the two groups 57.3% (55/96) vs 67.85% (19/28) respectively (two-tailed Fisher’s exact test, p=0.3842, Figure 1J). Likewise, 39.28% (11/28) of the *de novo* mutations had been reported in COSMIC, compared to 25% (24/96) of the germline mutations, which was also not statistically significant (two-tailed Fisher’s exact test, p=0.1568, Figure 1J). We next investigated similarities or differences between germline and *de novo* variants in terms of affected genes. Twelve genes carried both deleterious and non-deleterious germline SNVs, whereas in the *de novo* SNVs, the genes with deleterious mutations were unique and distinct from those with non-deleterious variants. Additionally, the germline and *de novo* SNVs had four genes in common (Figure 1K). Taken together, this shows that the germline and *de novo* SNVs differ in terms of potential functional impact. We then classified all the SNVs based on the cancer types to which the gene mutations typically associate. The results show that there is not an especially enriched mutational profile associating with specific types of cancer in germline nor *de novo* SNVs (Figure 1L). Lastly, we categorized the variants according to the gene’s function. Notably, *de novo* SNVs were predominantly affecting genes involved in transcriptional regulation and chromatin remodelling and were statistically significantly more often deleterious than their germline counterparts (p=0.0154, Fisher’s exact test, Figure 1M).

Mutations in cancer-related genes have been previously identified in hPSCs (Avior et al., 2021; Lezmi et al., 2024; Merkle et al., 2017, 2022). These studies used various approaches to screen hPSC lines, focusing on Tier 1 COSMIC-reported variants. In a seminal report, Merkle et al. highlighted the recurrent acquisition of *TP53* mutations in hPSCs (Merkle et al., 2017). Similarly, Avior et al. identified *TP53* mutations as the most frequent in the H1 and H9 lines, along with mutations in *EGFR, PATZ1*, and *CDK12* (Avior et al., 2021). In a more recent large-scale follow-up study, Merkle et al. found 382 Tier 1 cancer-associated mutations across 143 lines via whole genome sequencing, though they could not determine the mutation origins since only a single sample per hPSC line was tested (Merkle et al., 2022). Lezmi et al. conducted an in-depth analysis of mRNA sequencing data and reported that 25% of the 146 hPSC lines they studied carry cancer-associated mutations (Lezmi et al., 2024). In 70% of cases, they find the mutations to appear *de novo* in culture or during differentiation. In our work, the study of multiple passages of the same line has allowed us to establish with certainty if a variant appeared *de novo*, and when in time in culture this occurred, a data point that was missing in the previously published work. In our cohort, we find that eight of the *de novo* and fourteen of the germline SNVs are Tier 1 COSMIC mutations. 65% of our lines carried a Tier 1 COSMIC variant, 70% of them having acquired the variant in culture. When looking at which genes bear the mutations and the overlap with previous work, we also find *TP53* as most recurrent, but also *KMT2C* (appearing twice in our study and identified by Lezmi et al and Merkle et al). In our study, both *TP53* variants are COSMIC-reported variants, one decreases in frequency with time in culture and the second is homozygous, suggesting a loss of heterozygosity. Other genes in common with previous reports are *CREBBP, FAT1, PMS2, BRCA2* and *APC*, the last four appearing as germline variants in our study. *BCOR* mutations that have been reported in hiPSC (Rouhani et al., 2022) were not found in our study nor previous work (Avior et al., 2021; Lezmi et al., 2024; Merkle et al., 2022). Sporadic mutations in hPSCs can have significant phenotypic effects, limiting their utility in clinical applications. Amir et al. showed that hESCs with *TP53* mutations gained a selective advantage under stressful culture conditions and retained a higher percentage of cells expressing the pluripotency marker *OCT4* after differentiation, resulting in increased cell proliferation and survival rates (Amir et al., 2017). Further, Lezmi et al. showed that *TP53*-mutated hPSCs had decreased neural differentiation capacities (Lezmi et al., 2024).

In the last step of analysis, we integrated the CNV and SNV data, to test for associations between the two. In previously published work that tested for chromosomal abnormalities (Avior et al., 2019; Merkle et al., 2022), identifying a link between CNV and SNV proved challenging because of either the relatively lower resolution of e-karyotyping (Avior et al., 2021) or the lack of multiple passages of the same line (Merkle et al., 2022). In our study, it is important to bear in mind that we do not always have a perfectly maintained single lineage within the lines because some of the later passages of our lines are from historically frozen vials. Also, although some lines were maintained in culture continuously for years, they clearly drifted into genetically different sublines.

A first finding from integrating the CNV and SNV data is that the three germline variants that had frequencies that were at odds with being heterozygous or homozygous (indicated with a triangle in Figure 2) were in fact located in a duplicated chromosomal region. VUBe026 carried at the earliest passage tested (Passage 11) a duplication of 9p24.3p13.3. This region contains *CDKN2B* that carries a SNV with an allelic frequency of 0.7 as well as a deleterious and COSMIC-reported *DOCK8* mutation with an allelic frequency of 0.25. The duplication of 9p24.3p13.3 explains the increased frequency of the *CDKN2B* SNV, as the line was likely heterozygous at the start, and the region with the variant was duplicated. The frequency of the *DOCK8* mutation suggests that it may be a *de novo* mutation that occurred shortly after or concurrently with the gain of 9p24.3p13.3, and in linkage with the *CDKN2B* variant, explaining its frequency as a single copy in a triplicated locus. VUBe014 carries a *PMS2* variant with an allelic frequency of 0.63 by passage 88 and has simultaneously acquired an abnormal karyotype with a gain of the 7p arm, spanning the *PMS2* gene. A second and significant finding is that majority of *de novo* SNV appear to be following the acquisition of chromosomal abnormalities, especially those known to confer a growth advantage to the cells. Four of the 13 karyotypically normal samples carried a *de novo* SNV, in contrast to 17 of the 20 karyotypically abnormal passages (30.7% vs 85% respectively, p=0.0028, Fisher’s exact test). In all but three of the latter 17 instances, the abnormal karyotypes included gains of 20q11.21 and 1q, making it challenging to determine whether any of these SNVs individually confer a growth advantage to the cells or if they are merely passengers or complementary to the chromosomal abnormality. In VUBe024 and VUBe026 SNVs associated to CNVs other than the gain of 20q or 1q. VUB024 carried a small gain of 1q spanning genes with no obvious beneficial effect if duplicated as well as a low frequency deleterious variant in *ASLX1*. Both were lost in later passages, suggesting they did not confer a growth advantage to the cells. VUBe026 is a more notable exception, where a gain of a deleterious variant in *EP300* occurred between passages 11 and 35 and persists in different sublines that acquired an array of different chromosomal abnormalities. This suggests that this specific variant may be conferring an *in vitro* advantage to the cells. By comparison, Avior et al identified only one trisomy 17 concurrently with one of the TP53 mutations (Avior et al., 2019), but their method focussed on detecting trisomies of 1, 12, 17 and 20, and would not have detected any of the gains of 20q found in our study, and only one of gains of chromosome 1. In the large study by Merkle et al, half of the 14 SNVs they identified as with highest oncogenic potential were associated to aneuploidies (Merkle et al., 2022). In our study, five SNVs are not concurring with CNVs, two are COSMIC-reported mutations (c.15461G>A in KMT2D (VUBe002) and c.2926G>A in KMT2C (VUBe003)), two are potentially deleterious (c.5934_5935dup in *FAT3* (VUBe002) and c.238C>A in *CYSLTR2* (VUBe005)) and the last one is a non-deleterious variant (c.4071G>C in *MET* (VUBe004)). The variant observed in *CYSLTR2* is the sole abnormality acquired by the VUBe005 line at a latest passage tested, whereas the other variants disappeared in the later passage of the lines. It is possible that the mutations in the *KMT2D* and *FAT3* genes persisted in an alternative subline of VUBe002, as they were detected at passage 36, with the next and final passage tested being passage 353, corresponding to a prolonged period in culture. Whether variants in these genes can have a role in promoting *in vitro* growth advantage in hPSCs remains to be elucidated.

## CONCLUSIONS

Our comprehensive analysis of the hESC lines across multiple passages provides insights into the dynamics of genomic alterations during *in vitro* culture. We observed frequent emergence of both *de novo* CNVs and SNVs in cancer-related genes throughout culture, with mutation rates remaining stable over time, indicating that the higher mutational burden in later passages is cumulative due to prolonged culture rather than increased mutagenesis. Notably, the *de novo* SNVs often affected genes involved in transcriptional regulation and chromatin remodelling, with a higher proportion being deleterious and reported in COSMIC compared to germline SNVs. Integration of the CNV and SNV data revealed that CNVs that are not typically recurrent in hPSC cultures frequently emerge in association with known CNVs that confer a growth advantage to hPSCs, such as the gains of 1q and 20q11.21, and that many *de novo* SNVs appeared after the acquisition of these recurrent chromosomal abnormalities. These associations suggest two potential scenarios: either rare CNVs and the majority of the SNVs were passengers during the culture takeover of the recurrent CNVs or they potentially interact and enhance these CNVs. While functional experiments are necessary to fully understand their impact in regenerative medicine, it is rather reassuring that most of the low-passage and genetically balanced samples were devoid of de novo Tier 1 COSMIC mutations, ensuring their safety for use in research and therapeutic applications.

## SUPPLEMENTARY MATERIAL

**Supplementary Table 1**. Overview table of the genes mutated and type of variants: general, all lines together, overview of the detected mutations, variant load/frequency.

**Supplementary Table 2**. Table with the sequenced genes: genes panel.

**Supplementary Table 3**. Table showing the panel of 380 cancer-associated genes used in the analysis.

## MATERIALS AND METHODS

### Institutional Review Board Statement

For all parts of this study, the design and conduct complied with all relevant regulations regarding the use of human materials. Ethical review and approval were waived for this study because it involved work on existing in-house derived hESC lines.

### hESCs lines and cell culture

All hESC lines were derived and characterized as reported previously (Geens et al., 2009; Mateizel et al., 2006, 2010), details on the characterization of the lines can also be found at the open science framework (https://osf.io/esmz8/) and are registered in the EU hPSC registry (https://hpscreg.eu/). The cells were cryopreserved in freezing medium composed of 90% knock-out serum (Thermo Fisher Scientific) and 10% dimethyl sulfoxide (Sigma-Aldrich). In the past, our hESC lines were cultured on 0.1% gelatin-coated (Sigma-Aldrich, Schnelldorf, Germany) culture dishes containing mitomycin C inactivated CF1 mouse embryonic fibroblast (MEF) feeders, in hES medium culture medium supplemented with 20% KO-serum replacement (Mateizel et al., 2006). Cells were passaged by manual dissection of undifferentiated cell colonies. Currently, hESCs are cultured on tissue culture dishes coated with 10 µg/mL Biolaminin 521 (Biolamina®) and maintained in NutriStem hESC XF medium (NS medium; Biological Industries) with 100 U/mL penicillin/streptomycin (P/S) (Thermo Fisher Scientific). Cells are passaged as single cells using TrypLE Express (Thermo Fisher Scientific) and split at a ratio of 1:10 to 1:100 as needed at 70-90% confluence. Cells were fed daily with NutriStem hESC XF medium in a 37 °C incubator with 5% CO2. All cultures are monthly tested for the presence of mycoplasma. For this study, hESC that had been cryopreserved from MEF cultures, were thawed on Biolaminin-521 and NS medium and expanded for few passages to obtain sufficient cells to extract DNA for the analysis. The identity of all samples in this study was authenticated by fingerprinting on the same DNA sample used for sequencing. Supplementary table S1 indicates which lines were kept on MEF prior to their cryopreservation and thawing for this study. We studied a total of 33 samples collected across 10 hESC lines: VUBe001 (passages 66, 117, 285), VUBe002 (passages 6, 36, 353), VUBe003 (passages 16, 61, 98), VUBe004 (passages 16, 50, 111), VUBe005 (passages 39, 75, 89), VUBe007 (passages 26, 40, 88, 198, 208), VUBe014 (passages 20, 50, 88), VUBe019 (passages 26, 60, 79), VUBe024 (passages 25, 43, 75), and VUBe026 (passages 11, 35, 39, 54).

### Fingerprinting

DNA fingerprinting was done with the Devyser Complete v2 kit (Devyser). Briefly, a multiplex PCR was carried out which interrogates 32 STR markers on chromosomes 13, 18, 21, X and Y. Separation of the different amplicons was done on a Genetic Analyzer 3500 (ABI) and Genemapper v6 (Themo Fischer Scientific) was used for subsequent data interpretation.

### Whole-genome shallow sequencing

The genetic content of the hESCs was assessed through shallow whole-genome sequencing by the BRIGHTcore of UZ Brussels, Belgium, as previously described (Bayindir et al., 2015). 5 µl of purified DNA is processed using the KAPA HyperPlus kit (Roche Sequencing, CA, USA) according to manufacturer’s recommendations, with five modifications: (1) an enzymatic fragmentation for 45 min at 37°C to obtain DNA insert sizes of on average 200 bp, (2) the usage of 15 µM of our in-house designed UDI/UMI adapters (Integrated DNA Technologies, Coralville, IA, USA), (3) a 0,8x post-ligation AMPure bead cleanup, (4) a total of 6 PCR cycles are applied to get sufficient library and (5) a 1x post-PCR AMPure bead cleanup. The final library is quantified and qualified with resp. the Qubit 2.0 using the Promega Quantifluor ONE kit (Promega, WI, USA) and the AATI Fragment Analyzer (Agilent Technologies Inc., Santa Clara, CA, USA) using the DNF-474 High Sensitivity NGS Fragment Analysis Kit (Agilent Technologies Inc., CA, USA). The final library is diluted to 1,5 nM prior to denaturation for analysis on a NovaSeq S1 100 cycles run (Illumina Inc., CA, USA), generating on average 7 million 2×50bp reads. Following demultiplexing with bcl2fastq (v2.19.1.403) all reads are mapped to the human genome (UCSC b37) using BWA aln v.0.7.10. The aligned reads are sorted based on genomic coordinates using samtools sort (v0.1.19) and duplicates are removed with the Picard markduplicates tool (v.1.97). Following removal of blacklisted regions (in-house table), the coverage is calculated in bins of 50 kb with bedtools coverageBed (v2-2.25.0). Following GC correction, Z score, fold change and log2ratio calculation using in-house developed R scripts, the data is visualized in an in-house developed tool called BRIGHTCNV. Part of the data visualization in BRIGHTCNV is making use of JBrowse v1.0.1.

### Gene-panels for cancer-associated genes

Libraries were constructed on 150 ng of input DNA with the KAPA HyperPlus kit (Roche Sequencing, CA, USA) according to manufacturer’s recommendations, with three modifications: (1) an enzymatic fragmentation for 20 min at 37°C was used to obtain DNA insert sizes of on average 200 bp, (2) the usage of 15 µM of our in-house designed UDI/UMI adapters and (3) a total of 8 PCR cycles were applied to get sufficient library for target enrichment. Target enrichment was performed according to version 5.0 of the manufacturer’s instructions with a homebrew (STHT v3) KAPA HyperChoice probemix (Roche Sequencing, CA, USA), Pre-capture pooling was limited to max. 8 samples for a total of 1,2 µg of pooled library. In contrast to the instructions, the xGen Universal Blockers TS Mix (Integrated DNA Technologies, Coralville, IA, USA) replaced the sequence-specific blocking oligos and the final PCR was limited to 11 PCR cycles. Final libraries were qualified on the AATI Fragment Analyzer (Agilent Technologies Inc., Santa Clara, CA, USA), using the DNF-474 High Sensitivity NGS Fragment Analysis Kit (Agilent Technologies Inc., CA, USA) and quantified on the Qubit 2.0 with the Qubit dsDNA HS Assay Kit (Life Technologies, CA, USA). Per sample, a minimum of 14,5 million 2×100 bp reads were generated on the Illumina NovaSeq 6000 system (Illumina Inc., CA, USA), with the NovaSeq 6000 S4 Reagent Kit (200 cycles) kit. For this, 1 nM libraries were denatured according to manufacturer’s instruction.

The raw basecall files were converted to .fastq files with Illuminas bcl-convert algorithm. Reads were aligned to the human reference genome (hg19) with bwa (version 0.7.10-r789). Duplicate reads were marked with picard (version 1.97). Further post-processing of the aligned reads was done with the Genome Analysis Toolkit (GATK) (version 3.3). This post-processing consisted of realignment around insertions/deletions (indels) and base quality score recalibration. Quality control on the post-processed aligned reads was done with samtools flagstat(version 0.1-19) and picard HsMetrics(version 1.97). Those tools were used to investigate the total number of reads, the percentage of duplicate reads, the mean coverage on target and the percentage of on-target, near-target, and off-target bases). Variants were called using GATK Mutect2 (version 4) in tumor-only mode and annotated using annovar (version 2018Apr16). Variants with population frequency higher than 1% in gnomad, 1000g and esp6500 were filtered out. Furthermore, non-hotspot variants with an allele frequency below 3% were filtered out.

The called variants in VCF files were visualized using Basespace Variant Interpreter online tool (https://variantinterpreter.informatics.illumina.com). Recurrent variants defined as variants occurring in > 0.01% of samples sequenced over time at BrightCORE facility using the same gene panel were filtered out. A variant read count of ≥ 25 was used as cutoff to keep a variant in the analysis. After filtering the variants were manually inspected on IGV (https://igv.org). Any suspected genomic regions such as GC repeats, indels at microsatellite regions etc. were removed from analysis. The gene CDC27 was excluded from analysis due to high prevalence of pseudogenes (Kazemi-Sefat et al., 2021).

### Materials and availability statement

All VUB stem cell lines in this study are available upon request and after signing a material transfer agreement. Raw sequencing data of human samples is considered personal data by the General Data Protection Regulation of the European Union (Regulation (EU) 2016/679). The data can be obtained from the corresponding author upon reasonable request and after signing a Data Use Agreement. The source data for all figures is available as a supplementary table. The data supporting all figures in this paper can be found at the Open Science Framework repository (https://osf.io/y8tzh/).

## Acknowledgments

The authors wish to thank An Verloes and Brecht Ghesquiere for their technical and logistical support in the laboratory.

## Author Contributions

Conceptualization, C.S.; Methodology, D.A.D., M.S.G., N.K., A.H., M.R., C.M.D and Y.L.; Software, C.O., D.A.D. and M.S.G.; Validation, D.A.D, M.S.G. and C.O.; Formal Analysis, D.A.D, C.S. and M.S.G.; Investigation, D.A.D, C.S. and M.S.G.; Resources, C.S. and K.S.; Data Curation, D.A.D, M.S.G. and C.O.; Writing – Original Draft Preparation, D.A.D and C.S.; Writing – Review & Editing, D.A.D, K.S. and C.S.; Visualization, M.R., D.A.D. and C.S.; Supervision, C.S.; Project Administration, C.S.; Funding Acquisition, C.S. and K.S.

